# Cell twisting during desiccation reveals axial asymmetry in wall organization

**DOI:** 10.1101/2021.11.17.467786

**Authors:** Sedighe Keynia, Thomas C. Davis, Daniel B. Szymanski, Joseph A. Turner

## Abstract

Plant cell size and shape are tuned to their function and specified primarily by cellulose microfibril (CMF) patterning of the cell wall. *Arabidopsis thaliana* leaf trichomes are responsible for protecting plants against environmental elements and are unicellular structures that employ a highly anisotropic cellulose wall to extend and taper, generating pointed branches. During elongation, the mechanisms by which shifts in fiber orientation generate cells with predictable sizes and shapes are unknown. Specifically, the axisymmetric growth of trichome branches is often thought result from axisymmetric CMF patterning. Here, we analyzed the direction and degree of twist of branches after desiccation to reveal the presence of an asymmetric cell wall organization with a left-hand bias. CMF organization, quantified using computational modeling, suggests a limited reorientation of microfibrils during growth and maximum branch length limited by the wall axial stiffness. The model provides a mechanism for CMF asymmetry, which occurs after the branch bending stiffness becomes low enough that ambient bending affects the principal stresses. After this stage, the CMF synthesis results in a constant bending stiffness for longer branches. The resulting natural frequency of branches after a length of 200 μm falls within the range of the sounds associated with many insects.

**Statement of significance:** The growth of plant cell walls is governed by the direction of cellulose synthesis but the factors that influence the overall wall anisotropy are only partially understood. The twist of leaf trichome branches after desiccation reveals a left-handed asymmetry in cell wall organization even though the geometry is axisymmetric. This surprising behavior provides information about the directionality of cellulose synthesis control in plant cells.

## Introduction

The shape and growth pattern of a plant cell are determined by the organization of cellulose microfibrils (CMFs) (1–4), the major load-bearing components within cell walls, that are needed to constrain the relatively high (∼5-10 atm) hydrostatic turgor pressure. The remaining cell wall volume consists of a hydrated matrix of pectin, hemicellulose, and structural proteins (5, 6). According to the cellulose synthase constraint hypothesis (7, 8) cortical microtubules (CMTs) and associated proteins control the synthesis of CMFs, and CMT orientation has been correlated with cell wall stresses (9, 10). Thus, tracking of CMTs is often used to reveal information about CMF organization during growth, and the anisotropic mechanical properties that arise from their orientation.

According to the multinet growth hypothesis (11, 12), CMFs realign to the direction of maximum stress. Unfortunately, it is not possible to measure stress in cell walls directly, so computational models are needed to match measured load-displacement and/or strain data to infer mechanical behavior. Then, model stress patterns can be used to reveal relations between CMTs and CMFs. Such analyses for most plant tissues are complicated by cell-cell interactions because boundary conditions are challenging to estimate. For this reason, isolated, unicellular structures, such as the leaf trichomes studied here, provide valuable insight into the role of stress and cell wall heterogeneity for cell growth (13).

The organization of cell wall constituents controls the wall mechanical behavior. In particular, the orientation distribution of CMFs is critical to the determination of wall strain and the growth patterns under turgor pressure (6). However, when the turgor pressure is removed, the resulting deformation can reveal organization information that cannot be determined from the pressurized cell alone. A similar concept is used in biomedical engineering to study the material behavior of arteries (14, 15). An artery is a pressurized composite tube composed of smooth muscle cells, elastin, and collagen, each with different mechanical properties. Growth and remodeling in the loaded state of the artery produce stresses for the unloaded state. When a single axial cut is made in an unloaded artery, the cross-section opens at a specific angle because of residual stress in the artery wall. We exploit a similar concept through the removal of turgor pressure by desiccation; once the pressure is removed, the cell deformation reflects the stress state under pressure. Hence, by using a computational approach (cell wall deformation after dehydration and stress release) along with the experimental measurements, concepts about organization, such as the multi-net growth hypothesis, can be investigated.

In addition, recent studies of trichome dynamics (16, 17) used FE models to predict the natural frequencies associated with the different vibration modes. Their results were interpreted with respect to trichome functionality against herbivore insects and were analyzed relative to the sounds generated by the insects, but their predicted frequencies were higher than the range from many insects. However, their FE models were based on an isotropic material with mechanical properties that were constant during growth. Clearly, more accurate material information would improve the predicted vibration behavior.

In this article, desiccation of leaf trichome branches is used to quantify the cell wall mechanical properties using a computational model that reproduces the experimental deformations. The model is then used to understand changes in the axial stiffness, bending stiffness and natural frequencies of the branch during growth. The stiffnesses define the capacity of the branch to deformation for a specific loading direction and the natural frequency defines potential interactions against environmental threats. The results are expected to impact future studies focused on the role of genetics and/or environment on plant growth and development. Specifically, other important trichoblasts include cotton fibers (18, 19), for which small advances in modeling efforts could have a dramatic impact on the global textile industry. Computational approaches, when effectively integrated with advanced biological methods, may allow new crop varieties to be developed efficiently.

## Materials and Methods

### Microtubule imaging details

Plate grown Col-0 seedlings carrying a transgenic UBQ1p::TUB6-RFP construct were used for imaging trichomes on the first 5 leaves. Briefly, Z-stacks were taken using a 100X oil objective and a 561 nm excitation laser on a spinning disc confocal microscope for several overlapping sections of the trichome (as many necessary to capture the whole trichome branch from base to tip). Images were stitched together and processed using ImageJ, and microtubule orientation were analyzed from ROIs covering large portions of the trichomes using the ImageJ plugin OrientationJ.

### Microtubule angle quantification

Microtubule images were obtained on a spinning disk confocal microscope with a 100X 1.4 NA oil immersion objective, using a 561 nm solid-state laser as previously described (13). 11-13 DAG seedlings were whole mounted on chambered slides, and young leaves were imaged. An image tiling and montaging approach were taken to capture the entire cortical surface of a trichome branch. After brightness and contrast and Gaussian blur smoothing, background subtraction was performed using the rolling ball method with a 15-pixel radius. Images were sharpened using the unsharp mask function and oriented horizontally with the apex facing to the right. The ImageJ plugin OrientJ was used to quantify the dominant angle and coherency of the population of microtubules within each branch. Microtubule angle distributions were used to score each branch as either longitudinal/mixed, right-handed helical, or left-handed helical.

### Plant materials, growth conditions, trichome stage definition and desiccation

The Arabidopsis (*Arabidopsis thaliana*) Columbia and Ler ecotypes were used to observe trichome behavior under turgor pressure removal. For all experiments, a range of branch lengths from young trichomes (branch lengths less than 50 µm), to old trichomes (branch lengths longer than 300 μm), was examined during desiccation with time-lapse imaging. Seeds were stratified at 4 °C for 2 days and then grown in a growth chamber for 2-22 days under 12 h day length at 24 °C and relative humidity of 50–60 %. The goal was to study different trichome branches with many different lengths.

Complete removal of the turgor pressure was needed to measure the amount of twist with respect to branch length. The whole plant was kept in soil at room temperature for about one month without watering (this duration may be different if this protocol were to be executed in a different facility due to the difference in humidity levels and months of the year). During this time, the plants were exposed to day and night light conditions in which the plant is naturally grown. Using the slow desiccation process, the water inside the vacuoles dried out, leading to the collapse and twist of trichomes. For rapid desiccation, a vacuum was applied for 20 minutes which decreased the pressure from 1 to ∼0.05 mbar. Completely twisted samples (∼220) were selected and prepared for the observations and measurements. To identify the direction of twist only, removal of a fraction of the water content in the vacuole was sufficient because the direction of twist was evident even after partial dehydration. A plastic Petri dish (2 and 3.5 in) was prepared, and the bottom of the plate was covered with strips of double-sided tape. Each leaf was trimmed from its attachment to the petiole. If the first and second leaves were the last or the last visible leaves, the whole plant was trimmed from its attachment to the root to avoid any possible damage to the leaves. The empty spaces between the leaves were filled with desiccant beads (silica gel desiccants 2-1/4 × 1-1/2 inches) to expedite the moisture removal. If the leaves were not visible by the naked eye and only observable under a 10X microscope, less than one day (∼ 10 hours) was required to remove the turgor pressure and observe the direction of the twist. Typically, it took more than 10 days for the older and larger leaves or the trichomes farther away from the petiole to dry out and twist. Finally, after sufficient time (depending on the size of the leaf and the position of the trichome), the direction of twist was visible using a 20X lens of a laser scanning confocal microscope. Scanning electron microscopy (SEM) was only useful for this study for fully developed leaves and trichomes. At those stages of growth, enough space existed between trichomes for imaging. For small leaves, the SEM vacuum resulted in tangling of trichomes and bending of the leaf such that high-quality images were challenging to use for twist measurements. The amount of twist for each range of branch length was measured by selecting a point on one of the edges of the collapsed trichome branch and moving along the path of the edge and measuring the amount of sweep angle. For a better understanding, see Figs. S2a and b, which shows the method used to measure the length and amount of twist, respectively.

### Finite Element Model

A finite element model of the trichome branch response to desiccation was created using the commercial finite element software Abaqus 2019. Several factors, including geometry, material properties, and boundary conditions, play a role in the deformation pattern after turgor pressure removal. The model parameters were adjusted to match the amount and direction of twist of the branches at different stages of growth (i.e., length). A more detailed description of the FE model, including the sensitivity study for mechanical properties and estimates of shear strain, is included in the Supplementary Information. The trichome branch was modeled as a composite shell in which the branch diameter at the flank was constant with a radius of 8 µm (13). Because microtubules and cellulose synthase (CESA) complex have a similar alignment (13), the cell wall was modeled as an orthotropic laminate composite with single dominant fiber orientation. The range of this orientation was determined from the amount of twist. A layer of viscoelastic matrix, representative of the pectin in the wall, defined the time-dependent behavior of the cell wall using values based on pavement cell viscoelastic properties (20, 21). The matrix modulus and modulus of the orthotropic layer perpendicular to the cellulose fibers were assumed as 100 MPa (22) while the modulus of the orthotropic layer parallel to the fibers was assumed as 70 GPa (23–26). Assuming a Poisson’s ratio of 0.45 resulted in a shear modulus in all directions of 45 MPa. The wall matrix was assumed to constitute ∼36% of the cell wall with a density of 1000 kg/m (27), and the remaining volume was assumed to be occupied by cellulose (28), with a density of 1650 kg/m (25) which result in the whole cell wall density to be 1416 kg/m^3^, consistent with measurements (29) and an elasticity of ∼45 GPa in the direction of fibers and ∼100 MPa in perpendicular direction in case of fully aligned fibers, which is close to the range represented by Gibson (30). The trichome branch was modeled as a shell reservoir under constant hydrostatic turgor pressure with a pressure on the outer wall to represent atmospheric pressure (0.1 MPa). The turgor pressure in the model was increased from 0 to 0.6 MPa, held constant for ∼20s, then was removed with the same rate as the increasing rate, and the branch was then under atmospheric pressure. For a more accurate model to represent the large deformation observed after desiccation, a nonlinear analysis was implemented. Prior to our study, there were no suitable assumptions for the trichome branch mechanical properties that lead to the same twist after the removal of turgor pressure. Thus, the branch model was constructed based on the trichome branch shape, and the model properties were iterated to find the fiber orientation that reproduced the measured behavior.

## Results and Discussion

### Trichome Branch Twist Reveals Cell Wall Organization

Trichome branches (*A. thaliana* WT Col-0) were observed using a desiccation assay to quantify changes in their deformation (Fig. 1). Complete removal of turgor pressure took anywhere from a couple of hours to ∼30 days depending upon leaf age and the cell position relative to the petiole. A fast desiccation treatment (∼20 minutes; see Methods and Fig. S2) showed that the twist was not affected by possible reorientation of CMTs during the dehydration process. The geometry of the branch was symmetric about its long axis (i.e., axisymmetric) for all lengths (Fig. 1a). However, the observed twist after desiccation cannot occur unless the wall has an organization which is not axisymmetric. An image of an extensive collection of trichome branches (Fig. 1b, c) shows the consistency of the behavior after desiccation. More than 220 trichome branches were studied, and the amount and direction of their twist were measured as a function of branch length. Interestingly, there was a clear length dependence to the twisting which occurred only for branches longer than ∼100 μm. Branches shorter than this value (Fig. 1d) showed minimal twist suggesting an axisymmetric organization of the cell wall, but a right-handed twist was observed at this stage in a small number of branches (less than 5 %). On the other hand, longer branches (Fig. 1e-g) always had a clear left-handed twist. The branches in early stages (< 100 μm length) recovered their original shape after rehydration (Figs. 1h and i), which showed that the response was elastic and reversible. This behavior may indicate an efficient mechanism for shape recovery during/after acute water stress. The twist per unit length (see Supplemental Information) became a constant value of ∼0.82 °/μm (Fig. 1j) for branches longer than the transition length (∼100 μm), indicating that the wall organization was uniform. The maximum amount of twist per unit of length (Fig. 1k) increased from ∼0.6 °/μm to ∼1.2 °/μm when the mean branch length was about 175 μm and then dropped to ∼0.82 °/μm for branches longer than 300 μm.

**Figure 1.**
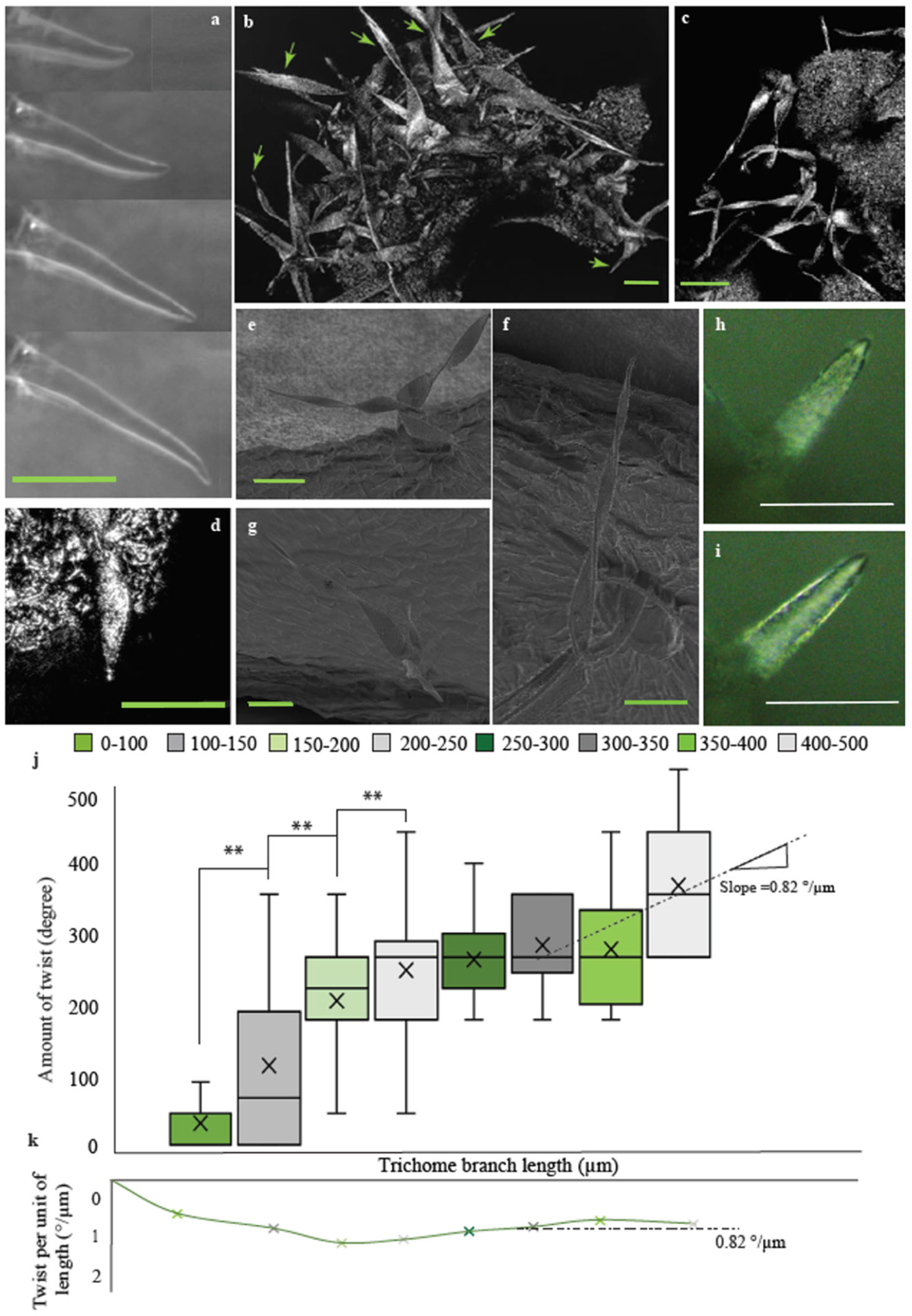
Twist of trichome branches during collapse after desiccation. (a) axisymmetric growth of leaf trichome branches (with permission from *Nature Plants* (13)). (b) and (c) are shows twisted trichome branches of *Arabidopsis* WT (Col) after desiccation. Arrows in (b) denote these ranches. (d) Right-handed twist of a short trichome branch observed in less than 5% of the branches and only when they were short. (e)-(g) Left-handed twist of trichome branch using SEM. (h) Trichome branch after dehydration. (i) The trichome branch in (h) after rehydration after ∼2 hours in water, which shows that the collapse of branches is elastic and recoverable; plastic deformation is not observed in the dehydration process. (j) Amount of twist and (k) twist per unit of length of desiccated branches for each range of length (in ° and °/ μm respectively), from measurements of more than 220 trichome branches. **p < 0.01. Scale bars = 40 µm in (a)-(i).

### Finite Element Model of a Trichome Branch

A finite element (FE) model was used to estimate the mechanical properties that would lead to the observed twist behavior. Although direct measurements of plant pavement cells have been made using nanoindentation (20, 21), atomic force microscopy (31, 32), and other related devices (33), such approaches are challenging with trichome branches because of their location above the plane of the leaf. For this reason, the initial cell wall material properties for the model were based on direct measurements from pavement cells (13). The FE model was then used parametrically to identify the ranges of material properties and organization (i.e., elastic moduli, fiber orientation) that were plausible relative to the deformations observed. Although there is an inherent non-uniqueness to this analysis, overall behavior can be quantified. To find a plausible set of material properties, different combinations of properties were assumed, and the results were compared with the deformations observed in desiccated trichome branches. The loading conditions for the model were defined using the turgor pressure, which was increased to a maximum value and held constant for a consistent amount of time which allowed the viscoelastic branch wall to relax fully. Then, the turgor pressure was removed so that the external atmospheric pressure was the only remaining applied force which collapsed the branch. The relaxation process allows residual stresses to develop in the cell wall. In arteries (14, 15), the growth-induced residual stresses and material properties determine the opening angle of the orthotropic structure after a single cut is made. By analogy, residual stresses in the cell wall develop during growth as part of the expansion process and their status is revealed by the desiccation. The trichome branches undergo large deformations after desiccation such that a nonlinear solver for large deformation behavior was used in the FE model.

The cell wall includes many constituents such as cellulose microfibrils, pectin, hemicellulose, and different proteins (5, 34). Computational models of plant cells must account for these constituents in some way. Their length scale relative to the cell size dictates either high-resolution and massive computational models with limited scope or models based on material homogenization, such as those used here, that allow extensive parametric studies to be conducted. Thus, the trichome branch cell wall was simulated as a homogenized composite shell with mechanical properties derived from cellulose and pectin in which the orientation of cellulose fibers was allowed to vary with respect to position (Fig. 2a; discussion of the sensitivity study is given in the Supporting Information). The properties of each element of the model were homogenized from the assumed composite organization to include both anisotropic and viscoelastic behavior. The axis of the transverse anisotropy, 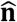, was defined such that 90° denoted alignment with the branch geometric symmetry axis and 0° with the radial direction. The viscoelastic properties, representative of the pectin-rich matrix, defined the time-dependent behavior of the cell wall (21). Cell wall thickness plays an essential role in the overall trend concerning the twist along the length of the trichome (Fig. 2b). Previous measurements (13) showed the presence of a thickness gradient from the tip towards the base, but those measurements focused on the early stages of growth and cell shape patterning. If a thickness gradient was used over the entire branch length, the FE models showed that the twist behavior was attenuated toward the base (Fig. 2b) suggesting a uniform thickness in longer branches along the majority of the length away from the tip (see Supporting Information). The layered, composite organization used here could include any number of layers and organization (Fig. 2a). We focused on a 3-layer composite to represent the ‘average’ wall behavior to simplify the quantification relative to the measurements. In this way, the trends with respect to length could be observed (Figs. 2c, d). As expected, the amount and direction of branch twist depended upon fiber orientation, whether left-handed or right-handed (Fig. 2d), with similar stress patterns. Based on the twist data, longer trichomes must have a higher percentage of fibers aligned more closely with the growth axis than shorter trichomes. However, if all fibers were oriented parallel with the geometric symmetry axis of the branch (axial direction), the model predicted wall collapse without twist (Fig. 2e), i.e., an axisymmetric collapse. Thus, the observed twist cannot be explained using fibers aligned only with the direction transverse to the geometric symmetry axis (circumferential) or aligned with it (axial).

**Figure 2.**
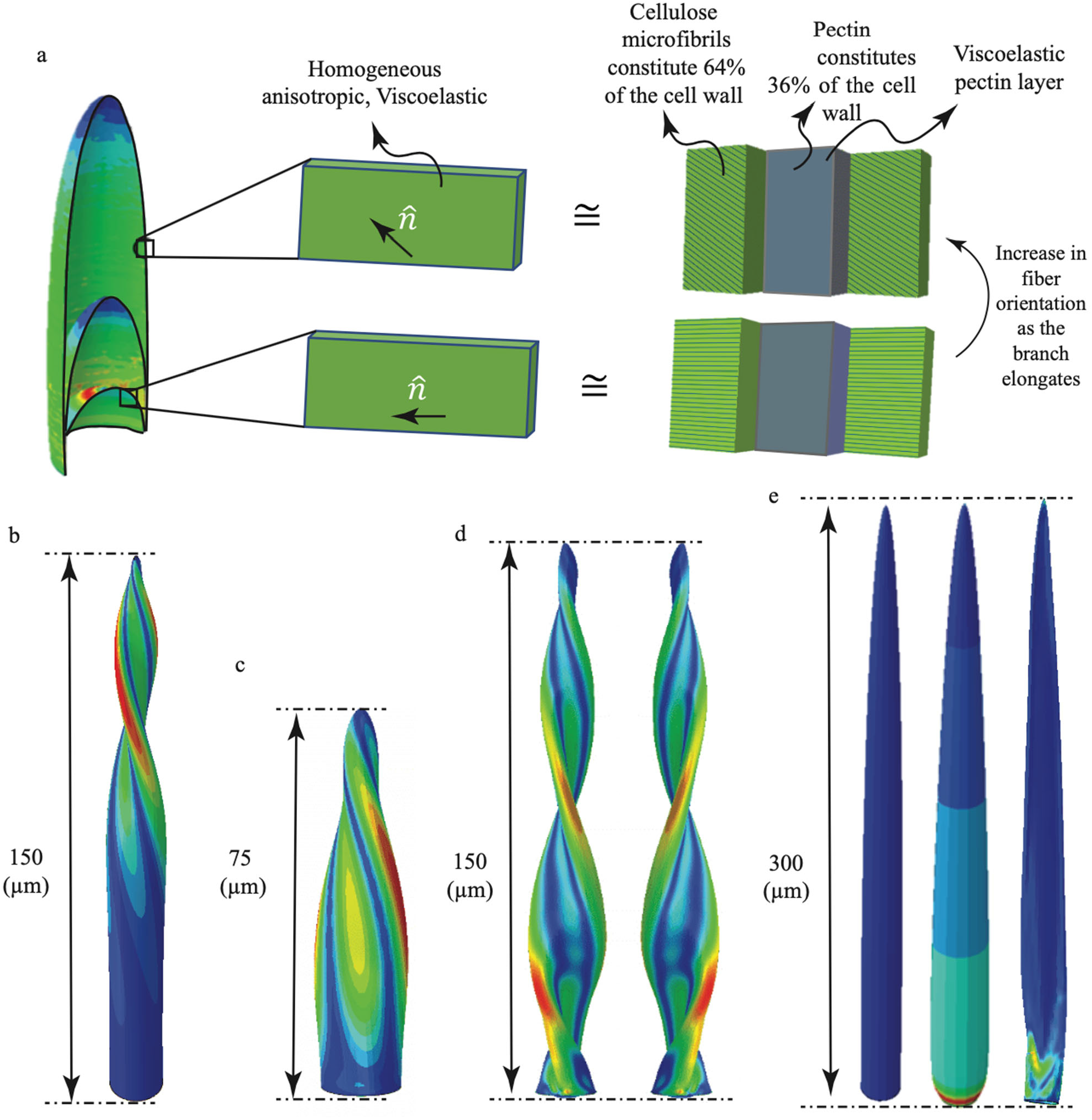
Finite element model of a trichome branch. (a) The organization of the cell wall is modeled as a composite shell of orthotropic and viscoelastic layers to represent the cellulose and pectin components, respectively, for different branch length. The composite of orthotropic and viscoelastic material is equivalent to a homogeneous anisotropic and viscoelastic material which is not available in FE software. (b) Effect of the thickness profile on branch twist; if the cell wall thickness increases continuously from the tip to the base the twist is confined to the tip only; a constant thickness of 200 nm for positions more than 50 nm from the tip leads to the same twist observed in experiments. (c) and (d) Cell wall von Mises stress of trichome branch model after removal of turgor pressure for branch length of 75 and 150 µm with dominant angle of -30 and -45 and 45-degrees, respectively. Effect of right-handed fiber orientation on the direction of twist is shown in (d). (e) FEM of trichome branch with all longitudinal fibers without pressure (left), swelled under constant pressure without any longitudinal strain (middle) and star-shaped collapse after removing the pressure (right).

### Microtubule Handedness is Consistent with Twist

The handedness of the twist of longer branches was clearly biased in the left-hand direction and highlighted the asymmetry in the cell wall that must be present. The angle of microtubule orientation can be used as an additional validation of the model because cortical microtubules (CMTs) and cellulose synthase (CESA) complexes are highly correlated in trichome branches (13). Live-cell microtubule (MT) imaging of entire branches (Fig. 3a-d) revealed clear patterns of cellular scale cortical microtubule organization. As previously reported (35), shorter branches had more transversely-aligned MTs while the orientation showed a directionality for longer branches. A bias toward left-handed asymmetry in trichome branch MT networks had been reported (36), but those measurements were limited with respect to trichome branch length and the fraction of the branch surface that was analyzed. We conducted high-resolution live-cell imaging of the cortical microtubule network of half of the branch cortex along its full length by tiling the individual images (Fig. 3a). The microtubule array usually had a consistent organization along the branch length, and mean MT orientation was quantified as a function of branch length. Figures 3e-f show a distinct change in organization from transverse in cells less than ∼100 μm to left-handed helical. Clearly, the MT patterning as a pathway for CMF synthesis exhibits a bias that matches the branch twist behavior that is dominated towards the left-handed direction. The consistency of trichome twist direction and handedness of MTs showed a preferred asymmetry for branch wall organization similar to the crystal texture of other systems in nature, such as giant barnacle shells (37).

**Figure 3.**
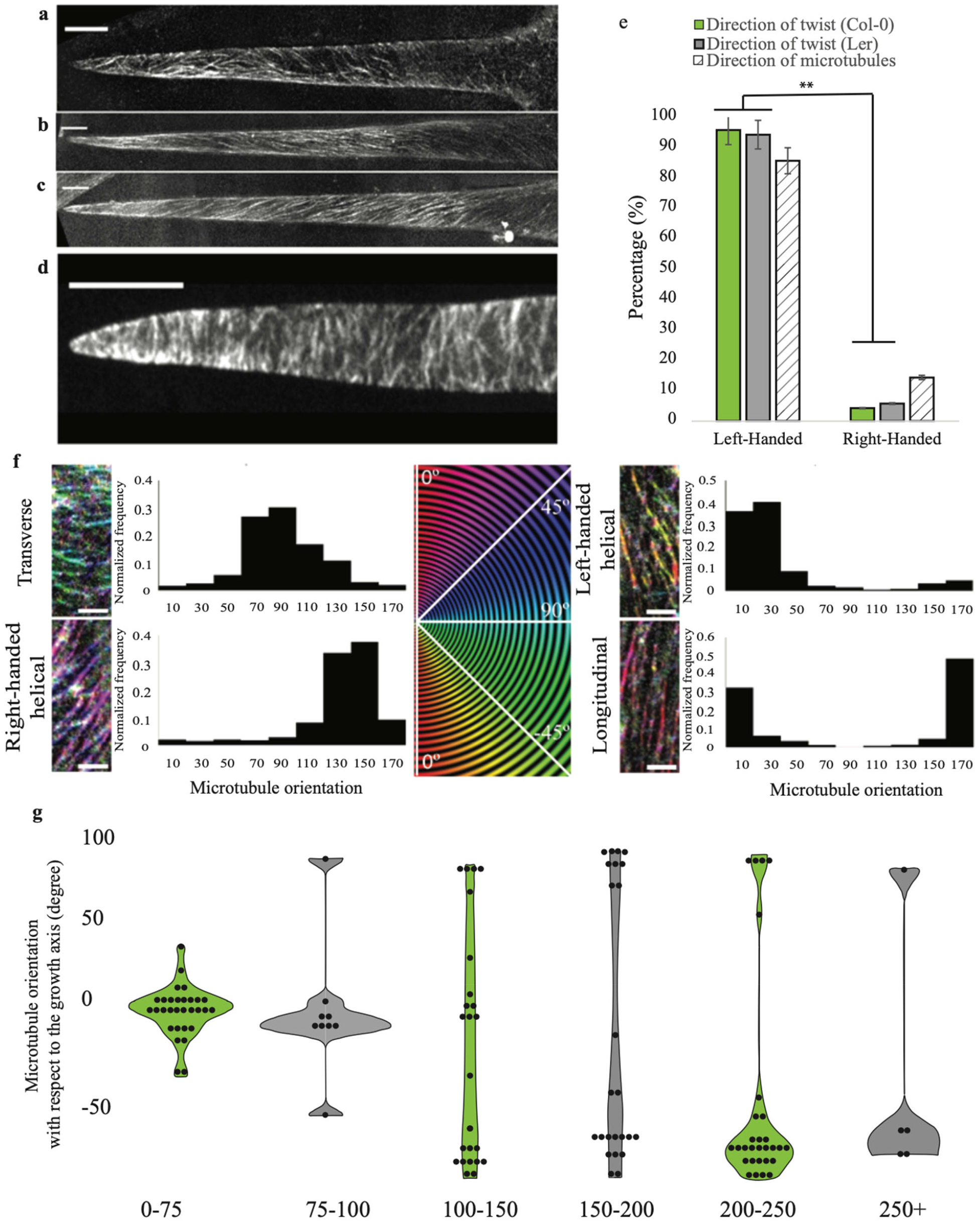
Microtubule alignment and degree of handedness at different stages of growth. (a)-(d) Cortical microtubules at the plasma membrane aligned in, right-handed direction (a), longitudinal direction (b), left-handed direction (c) and transverse direction (d). (e) Comparison of the handedness twist of desiccated branches in Col-0, Ler and microtubule orientation measurements. **P < 0.01. (f) Qualitative categorization of global alignment of branches. (g) Dominant angle of the MT alignment relative to the primary growth axis.

### Branch Stiffness and Natural Frequency

During branch growth, the newly synthesized CMFs change direction. The overall organization will affect the axial stiffness of the branch, defined as a weighted average of the material stiffness in the growth direction over the cross-sectional area of the wall (calculations provided in Supporting Information). Fig. 4 shows the computed axial stiffness as a function of branch length for the Col trichome branches based on the models that matched the twist data. The stiffness initially decreases as a function of branch length due to the dominance of transverse CMFs and corresponding lack of twist in desiccated branches. In longer branches, the axial stiffness increases because the CMFs are aligned more closely with the growth axis. The material displacement and stress at the base of the branch will be affected by the axial stiffness (i.e., higher stiffness will lead to lower displacement for the same turgor pressure). Growth stops after the computed axial stiffness approaches the critical value of ∼110 N/m (symbol indicated with an A in Fig. 4).

**Figure 4.**
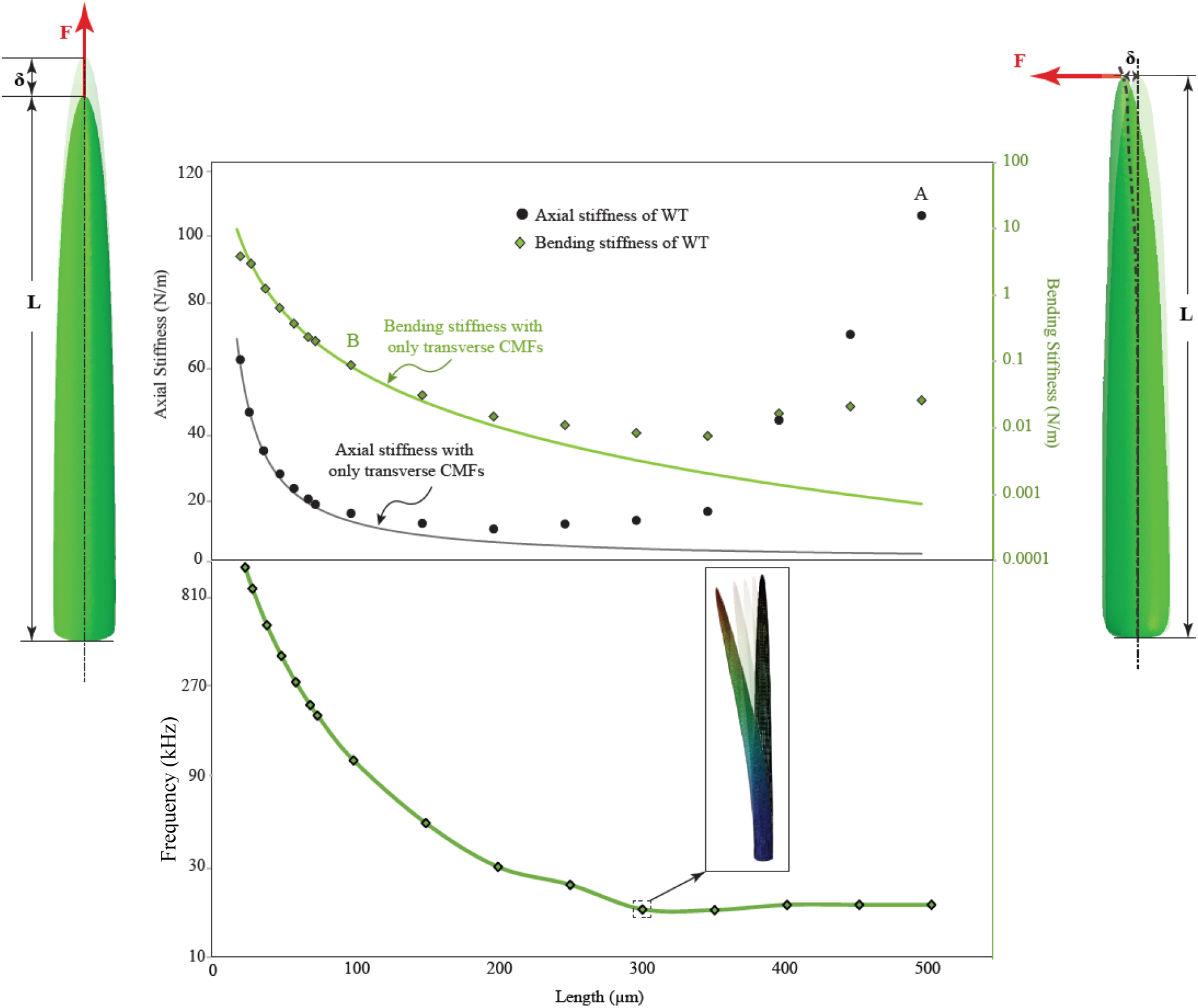
Influence of cellulose microfibril (CMF) orientation on bending stiffness and axial stiffness and natural frequency of the first mode. Predicted axial and bending stiffness of trichome branches with different lengths based on model values needed to match twist data. Axial stiffness (see Supplemental Information) converges to about 100 N/m at point A, which suggests that growth stops when the value of axial stiffness is reached. Bending stiffness for WT trichome branches shows an initial reduction but then a stabilization as a function of length. After a length of about 100 μm, when the bending stiffness reaches the minimum possible value, the CMF deposition is no longer transverse to the growth axis which stabilizes the influence of bending even for longer branches. Lowest frequency at which deformation occurs (first mode) after the length of ∼300 μm converges to ∼20 kHz with the change of branch length and cell wall material properties.

The mechanism for the change in CMF orientation observed after the transition length of 100 μm was examined using the model. An axisymmetric organization of CMFs in the wall, whether transverse to the geometric symmetry axis or aligned with it, would result in wall stresses that also reflect this axisymmetry. In other words, the principal stresses would align with the axial and transverse directions during the early stages of growth. Such an axisymmetric wall stress or strain could not induce an asymmetric CMF synthesis, nor could cause a passive realignment of existing CMFs. However, as the branch extends beyond a threshold value, the length itself could provide a mechanism to break the symmetry. The axial stiffness of the branch (Supporting Information) scales linearly with the axial modulus and inversely with length (∼*L*^-1^). The bending stiffness also scales linearly with axial modulus, but inversely with the ***cube*** of the length (∼*L*^-3^). In other words, as the branch grows with constant properties defined by the transverse CMF alignment, it becomes more susceptible to bending motion that would shift the direction of maximum principal stress in the cell wall and break the symmetry. This shift, in turn, could influence CMTs and the angle of cellulose fiber synthesis. Results from the Col FE model (solid green line in Fig. 4), based on an assumption of only transverse CMFs, show the trend for both axial stiffness and bending stiffness with respect to branch length. The model predicts that a branch of length ∼100 μm would have a bending stiffness of ∼0.08 N/m (symbol indicated with B in Fig. 4). Remarkably, models created to match the measured twist data predict a bending stiffness for longer branches that plateau to a constant value. Ambient environmental variations, such as thermal noise, could strongly influence such a compliant structure. A similar type of thermal noise is often exploited to calibrate atomic force microscope (AFM) cantilevers (38) that can be much stiffer (∼40 N/m), but more resonant. The estimated bending displacements of 10s to 100s of nanometers would be enhanced at the branch tip as tapering occurs due to the reduced cross-section. After the synthesis of oriented CMFs is initiated, the subsequent growth would never return to the condition of axisymmetric synthesis, and the principal stresses would remain unaligned with the geometric symmetry axis. At subsequent stages of growth, the shear stress could also cause some degree of passive reorientation of CMFs (11).

Finally, the first natural frequency of trichome branch bending was calculated based on the model values that matched the twist behavior (Fig. 4). The results show a convergence to a constant value of ∼20 kHz as the branch reaches the length of ∼300 μm. The vibration frequencies of leaf trichomes have been shown to correlate with protection of the leaves against insects such as caterpillars (16, 17). Here, the branch bending frequency converges to a value within the frequency band of several insects including caterpillars (16, 17), crickets (39), katydid (40) and cicadas (41).

## Conclusion

In this article, we have presented a predictive computational model that replicates the mechanical twist of desiccated trichome branches to provide insight into the organization of constituents within the branch cell wall that change during growth. The observed behavior cannot be the result of CMT reorientation or changes due to cell wall decomposition during the desiccation process because the twist was present even for rapid desiccation. The presence of an asymmetric organization is hidden by the axisymmetric geometry of the pressurized branch. The FE model was used to quantify the observations of the branch twist that revealed the changes that occurred during growth, and the MT imaging validated the wall organization necessary for twist. These observations do not support strong passive realignment of CMFs, although it may be present to some extent after the transition point of ∼100 μm at which time CMF synthesis asymmetry initiates. Shorter branches have only transversely oriented CMFs and such an organization would have principal stresses aligned with the axial and transverse directions. The length of the transition of CMF orientation corresponds with a predicted branch bending stiffness, based on models that match the measured twist, would allow ambient noise of the plant and environment to induce bending displacements of the branch. A combination of a bending motion and the disappearance of tip-localized cytoskeletal regulators (13, 42) may explain the length associated with the symmetry-breaking event that dramatically alters the CMF synthesis pattern. The degree of twist and direction of twist are controlled by cytoskeletal (43) and cell wall biosynthesis (44) systems that have poorly understood effects on cell wall patterning and morphogenesis. The approach described here provides a reliable strategy to integrate cell growth and desiccation phenotypes with biomechanics and their genetic control.

The results presented also suggest a limit to the hypothesis regarding passive reorientation. The multinet growth hypothesis suggests that cell wall elongation or expansion modifies the organization of CMFs that are exposed to shear stress in the cell, potentially resulting in passive reorientation toward the growth direction (11, 32). For a longitudinal strain of 14 %, the average orientation of wall polymer chains of onion was observed to rotate along the stretch direction by ∼5.3 ° (45). However, this tilt and realignment of CMFs have not been observed in all cell types. For example, live-cell imaging of *Arabidopsis* was used to analyze cellulose reorientation during root cell growth in WT seedlings (46). No evidence of reorientation was detected in a mutant, *procuste1-1* (*prc1-1*) with a partial defect in cellulose synthesis. Complete reorientation of all microfibrils to the symmetry direction would not support the twist measured here. After the left-handed deposition initiates, additional elongation of the branch would be accompanied by some reorientation of CMFs, which would enhance the axial stiffness of the branch. This behavior is thought to decrease the net growth rate of the cell, and eventually cease the growth process (46). By observing the growth of the WT *Arabidopsis* trichome branch, it is clear that the growth rate is almost constant during the first 50 µm. This rate increases for some periods of growth but never decreases (13). Thus, passive realignment cannot be the only control of the observed CMF orientation. Analysis of helical growth from *Arabidopsis* mutants such as the *tor2* or *tor2 zwichel* double mutant indicates that microtubules play a significant role in the helical phenotype of leaf trichomes (47). *Arabidopsis* mutants growing with a right-handed twist have been reported to have cortical microtubules that are oriented around the cell in left-handed helices and vice versa (48). The relationship between MTs and the direction of cell growth is usually explained by assuming that MTs control the deposition of load-bearing cellulose microfibrils of the cell wall, so the cells elongate or expand in a direction perpendicular to the deposited cellulose (13). Furutani et al. (47) used this model to explain helical growth in spiral mutants for which expansion perpendicular to cellulose fibers that are synthesized in a left-handed direction would result in right-handed growth and vice versa. Others explain the behavior as a helix that unwraps in the direction opposite to the direction of stretch so that a wall generated with a left-handed helical pitch would unwind with a right-handed twist as the cell elongates (49). The results here suggest that the helical growth observed in mutant trichome branches (50) cannot be the result of MT handedness alone because the handedness of synthesis in the WT gives rise to a straight branch.

Our results firmly establish that removal of turgor pressure can reveal salient information about cell wall composition, which can improve the interpretation of genetic and environmental effects on plant growth. Desiccation of cells has been used by others recently (51) to test hypotheses about changes in cell wall organization during growth. They showed that dehydration of cotyledons led to wrinkling of the periclinal cell walls while anticlinal walls were affected much less. If more accurate information about cell wall mechanical properties can be determined by means of desiccation, imaging, and computational models during growth, then more accurate predictions can be made regarding overall static and dynamic behavior. Previous models of trichome vibrations (16, 17), predicted frequencies above the range of many insects (Libersat et al., 1994; Pollack and Imaizumi, 1999; Bennet-Clark and Young, 1994), but those models were based on a thicker cell wall (1.5-6 μm) (52), isotropic properties, and lower cell wall stiffness (0.6-4.7 GPa) (53, 54). They also assumed that the properties were constant during growth. Our results based on the twist observations as a function of growth show that the frequencies would plateau when the branch reaches full maturity with a frequency closer to that of insects. Further research and multiscale models are needed to understand the role of mechanical stress on individual cell wall constituents to predict growth mechanics more accurately.

## Supporting Information

Supporting Information available on-line.

Supporting figures: Figure S1, Figure S2, Figure S3, Figure S4, Figure S5, Figure S6

Supporting tables: Table S1. Table S2.

Supporting animations: Movie.S1.

## Acknowledgments

We thank to the Purdue Life Science Microscopy Facility for their expert assistance. Thanks to the Nano-Engineering Research Core Facility (NERCF) for their high-performance microscopy and we acknowledge the Holland Computing Center (HCC). Both NERCF and HCC are supported in part by the Nebraska Research Initiative. This research was supported by NSF Grant No. IOS-1715444 to D.B.S. and J.A.T. The authors declare no conflict of interest.

## Author Contributions

All authors participated in the experimental design, data collection, data analysis and manuscript preparation. J.A.T. conceived the project. J.A.T. and S.K. developed the finite element models. S.K. conducted the branch imaging and data processing. T.D. performed the MT measurements.

